# Re-evaluation of some popular CRAC channel inhibitors – structurally similar but mechanistically different?

**DOI:** 10.1101/2024.09.26.615129

**Authors:** Yasin Gökçe, Sinayat Mahzabeen, Joshua Ghoorahoo, Matthew Thomas Harper, Nazmi Yaras, Taufiq Rahman

## Abstract

Ca^2+^-release activated Ca^2+^ (CRAC) channels mediate the store-operated Ca^2+^ entry (SOCE) that is used almost by all cells to replenish depleted intracellular Ca^2+^ stores. Aberrant CRAC channel functions have been implicated in various diseases including autoimmune diseases and as such these channels and the resultant pathway (SOCE) have been intensively studied by academic and pharmaceutical scientists over decades. To date, several small molecule inhibitors and few antibodies have been developed against these ion channels. Some small molecule inhibitors of the CRAC channels, namely BTP2, Synta66, Pyr6 and GSK-7975A, are widely used as tool compounds in the field but their mode of action remains unclear. In this study, we used whole cell electrophysiology supported by in silico modelling to propose likely binding site and binding mode of these four popular CRAC channel inhibitors.

## Introduction

Ca^2+^-release activated Ca^2+^ (CRAC) channels are ubiquitous and highly Ca^2+^-selective ion channels that mediate store-operated Ca^2+^ entry (SOCE) following physiological or pharmacological depletion of intracellular Ca^2+^ stores [1]. Besides this unusual mode of activation requiring internal Ca^2+^ store depletion, their uniqueness as ion channels also stems from the fact that they form on demand in foci where the endoplasmic reticular (ER) membrane comes in close proximity to the plasma membrane (i.e. the ER-PM junctions). A functional CRAC channel consists of the ER membrane-resident Stim proteins (serving as the Ca^2+^ sensor of the ER lumen) interacting with the plasma membrane-localised Orai proteins (forming the pore) following store depletion[2]. The Ca^2+^ influx mediated by CRAC channels (i.e. SOCE) not only replenishes the depleted internal stores but also manifests enough spatio-temporal complexities to regulate specific cellular functions including gene transcription, differentiation, proliferation, migration and exocytosis [1, 2].

Although initially thought to be a feature of non-excitable cells only, it is now clear that CRAC channels (thus SOCE) are present almost in all cells from both the central and peripheral nervous system. However, not all cell types rely on them to the same extent. Many immune cells, notably including T lymphocytes, manifest prominent SOCE and properly-functioning CRAC channels are indispensable for their proliferation and production of cytokines [1]. Consequently, excessive SOCE due to aberrant activity of CRAC channels (often because of overexpression of Stim and/or Orai proteins) seems to be a major pathophysiological feature underlying various autoimmune and inflammatory diseases [3, 4]. There are now pharmaceutical companies (including CalciMedica, Rhizen Pharmacueticals, Vivreon Biosciences, ChemiCare) with exclusive or major focus on targeting CRAC channels against a number of such diseases (e.g., acute pancreatitis, rheumatoid arthritis, psoriasis, asthma, some forms of cancer). In addition, some extended areas (e.g., stroke, acute renal injury, chronic pain, vascular remodelling, arterial thrombosis) that may benefit from inhibiting CRAC channels are also being actively explored [4].

SOCE has been extensively studied since its first description in the early 1990s [5, 6], contemporaneously spurring active campaigns to find pharmacological modulators of this pathway to aid basic and translational research [3, 4]. However, as the bona fide components of the CRAC channels i.e. Stim and Orai proteins only became known much later in 2005-2006 [7–11], almost all early small molecule inhibitors against SOCE were found or developed through phenotypic screening involving functional (e.g. Ca^2+^ imaging and/or whole-cell electrophysiology) and/or microscopy (e.g. nuclear translocation of NFAT) assays. Such methods yielded several popular SOCE-inhibitory chemical scaffolds such as the imidazoles (e.g. SKF-96365 and related antifungal agents), the boronates (e.g. 2-APB and its derivatives) and the bis(trifluoromethyl) pyrazoles. The latter class is represented by BTP2 (also known as YM-58483), which is a widely-used SOCE inhibitor and which also inspired subsequent development of other small molecule inhibitors of SOCE such as Synta 66, GSK 7975A and Pyr6 that are also frequently used to reduce SOCE[3]. However, the precise mechanism of action of the majority of these inhibitors remains largely unclear whilst for a few of them, there have been some suggestions about their putative binding sites on the Orai protein [4, 12–14].. Indeed, although BTP2 is widely used as a SOCE inhibitor, there appears to be wide variations in its efficacy, potency and kinetics of action across studies and cell types[15, 16]. Studies have indicated that BTP2 and some related compounds affect Orai1 function without affecting its coupling to Stim proteins [12, 16] but their precise mode(s) of recognition by Orai proteins remain hitherto unclear. In this work, we primarily used whole-cell electrophysiology of RBL-1 cells measuring *I*_CRAC_ (the characteristic whole-cell current representing SOCE/active CRAC channels) [1] to find out whether four widely-used CRAC channel inhibitors namely BTP2, Pyr6, Synta66 and GSK-7975A act intracellularly or extracellularly. Despite having comparable physico-chemical properties (**Table 1**), we reveal significant differences in their sites of action with respect CRAC channels in the plasma membrane. We then complemented our initial findings with *in silico* modelling of the possible binding modes of these popular tool compounds.

## Materials and Methods

### Cell Culture

Rat basophilic leukemia-1 (RBL-1, ATCC: CRL-1378^™^) cells were grown in Dulbecco’s modified Eagle’s medium (DMEM) supplemented with 10% (v/v) fetal calf serum, 2 mM L-glutamine and 1% (v/v) penicillin/streptomycin, as previously described [16, 17].

### Patch Clamp Electrophysiology

Native CRAC currents (*I*_CRAC_) were recorded from RBL-1 cells at the room temperature (25 °C) using the patch-clamp technique in its whole-cell configuration. Preparation of the patch pipettes, the composition of the internal (pipette) and extracellular (bath) solutions and recording protocol (including leak subtraction) were as described previously [16]. Briefly, currents were acquired through Axopatch 200B and Digidata 1440 A (Molecular Devices) in response to voltage ramps (−100 to +100 mV in 150 ms) applied every 2 s; data were digitized at 10 kHz and filtered at 1 kHz and pCLAMP 10 suite (Molecular Devices) was used for data acquisition and analysis. Peak *I*_CRAC_ amplitude (pA) was normalized to cell size represented by the capacitance (pF) and expressed as current densities (pA/pF). For experimental evaluation of the CRAC channel inhibitors, cells were pre-treated with blockers for ∼20 min prior to recording.

### Molecular Modelling and Docking

Homology modelling of rat Orai 1 (rOrai1, Uniprot ID: Q5M848, CRCM1_RAT) was performed through SwissModel server (https://swissmodel.expasy.org/), using published structure of Drosophila Orai (DmOrai, PDB ID: 4HKR) as the template. First 51 residues of rOrai1 sequence seems to be intrinsically disordered as predicted by the Alphafold, (https://alphafold.ebi.ac.uk/entry/Q5M848 and **Supplementary Fig 1**); therefore we excluded this portion from the sequence of rOrai1 before modelling. Initially, a dimeric form of rOrai1 model was subjected refinement through the GalaxyRefine2 server [18] and finally it was assembled into a hexamer based on the published hexameric DmOrai structure (PDB ID: 4HKR).

The 3D structures of all the four CRAC channel inhibitors used in this study were obtained from PubChem and their putative binding poses were predicted through *in silico* docking as described previously [19, 20]. Briefly, for each molecule, an initial blind docking was performed in three independent runs using AutoDock Vina version 1.2.5 with a GRID box encompassing the entire rOrai1 hexamer. The best ranked and reproducible pose of each molecule from such blind docking was then used as the reference pose for subsequent pose refinement through focused docking (10 independent runs for each molecule) using GOLD version 5.3 suite (CCDC, Cambridge) using the default ChemPLP score. The most reproducible poses amongst the highest ranked ones were considered as the most plausible binding mode for each molecule. For each docked complex, the 3D ligand interaction profiles were deduced through the Protein-Ligand Interaction Profiler (PLIP). All relevant figures were generated in PyMOL Molecular Graphics System (version 1.8.2.0 Open-Source, Schrödinger, LLC) whilst ligand interaction figures were obtained from PLIP web tool (https://plip-tool.biotec.tu-dresden.de/plip-web/plip/index). Where relevant, we cited hOrai1 (Uniprot ID: Q96D31)-specific amino acid residues that correspond to specific rOrai1 amino acids.

### Statistical analysis

Results were expressed as means ± SEM for n independent experiments done at least in triplicates. Statistical comparisons of the mean values were performed using GraphPad Prism 8 (GraphPad software Inc., CA, USA). Unpaired, Student’s t-test was used to compare the groups with respect to the control group. P values<0.05 was considered to be significant.

## Results and Discussion

### Evaluation of the effects of chosen inhibitors on *I*_CRAC_

We first began our study with BTP2 which is a protypical and most widely-used SOCE inhibitor belonging to the *bis*(trifluoromethyl)pyrazoles or the BTP class [3]. The latter was first identified in 2000 as inhibitors of interleukin-2 (IL-2) synthesis by scientists from Abbot Laboratories in a high throughput reporter (luciferase) gene-based screen of Jurkat T cells with an overall goal to find better, non-macrocyclic immunosuppressants [21]. Members of the BTP series were found to manifest good structure-activity relationship in retarding nuclear translocation of NFAT and thus reducing IL-2 production, although little was then known about the underlying molecular mechanism [21]. A specific member belonging to the BTPs, namely BTP2 (also known as YM-58483) was later found to be inhibitor of SOCE [15] and CRAC channels [14].

As can be seen in **Fig 1**, BTP2 when added to the extracellular (bath) solution at 10μM, it significantly suppressed *I*_CRAC_.

**Fig 1.**
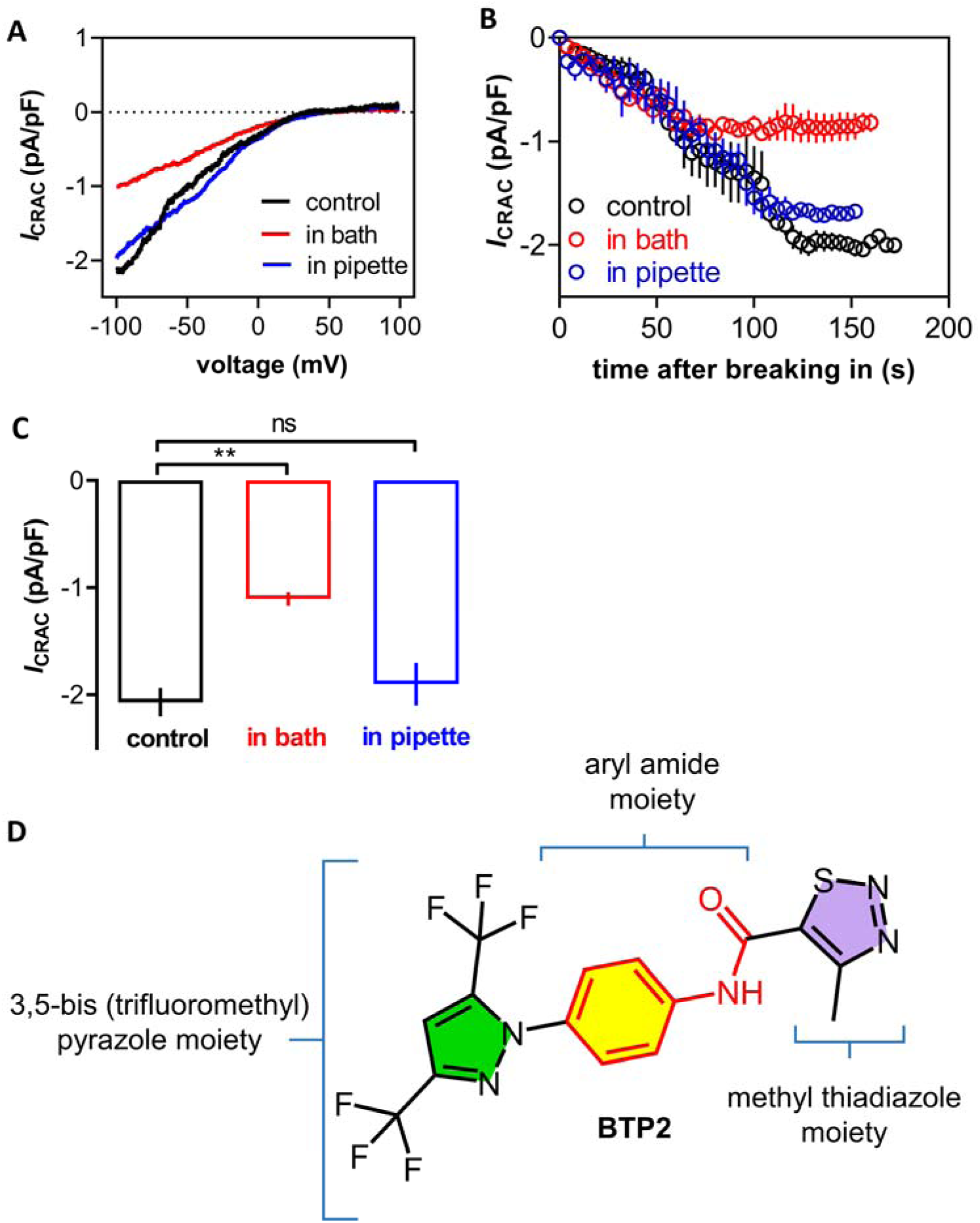
Effects of BTP2 on CRAC channel function in RBL-1 cells. Prior to recording of the whole-cell current representing CRAC channel function (*I*_CRAC_), RBL-1 cells were pre-treated with BTP2 (10μM) for ∼20min added to the bath solution or included in the pipette solution whilst control cells were treated with DMSO only. **A)** Typical current-voltage (I-V) relationships of *I*_CRAC_ (pA) evoked by voltage ramps from −100mv to +100mv, normalized to cell size (pF) under different experimental conditions. **B**) Representative peak *I*_CRAC_ size (as current density, pA/pF) vs time plots (each data point as mean ± SEM) from whole-cell recording of at least four different cells (n=4) in each condition are shown. **C)** Bar graphs (mean ± S.E.M) showing current densities at −80 mV holding potential under each condition. The statistical comparisons with respect to the control were done using Student’s unpaired t-test, **P<0.01. **D**) 2D chemical structure of BTP2, highlighting different sub-structural features.

However, at the same concentration included within the intracellular (pipette) solution, it seemingly caused a modest decrease in peak *I*_CRAC_ size which was not statistically significant. Thus, our result argues in favour of an extracellular mechanism of action of BTP2 against CRAC channels. Interestingly when first characterized against SOCE in Jurkat T cells, BTP2 was found to inhibit SOCE upon acute challenge when the Ca^2+^ entry plateaued and at this acute mode of challenge, it was as effective as it was with prolonged (10min) pre-incubation of the cells. The authors therefore suggested some sort of direct interaction with CRAC channels at surface level as possible mechanism of action of BTP2 [15].

Although results from a contemporaneous study [14] using similar cells differed significantly in finding the potency and the kinetics of SOCE blockade upon acute challenge of BTP2, it did confirm extracellular site of action of this CRAC channel inhibitor since its inclusion within the intracellular solution (at 1μM) did not affect *I*_CRAC_. Our finding with RBL-1 cells about BTP2’s site of action therefore strongly aligns with these observations in Jurkat T and related cells [14, 15].

Next, we sought out to test Pyr6 which is another notable SOCE inhibitor belonging to the same chemical class (i.e. BTP) [3]. Chemically, this molecule is effectively BTP2 with its methylthiadiazole moiety replaced by a fluoropyridine moiety (**Fig 2**).

**Fig 2.**
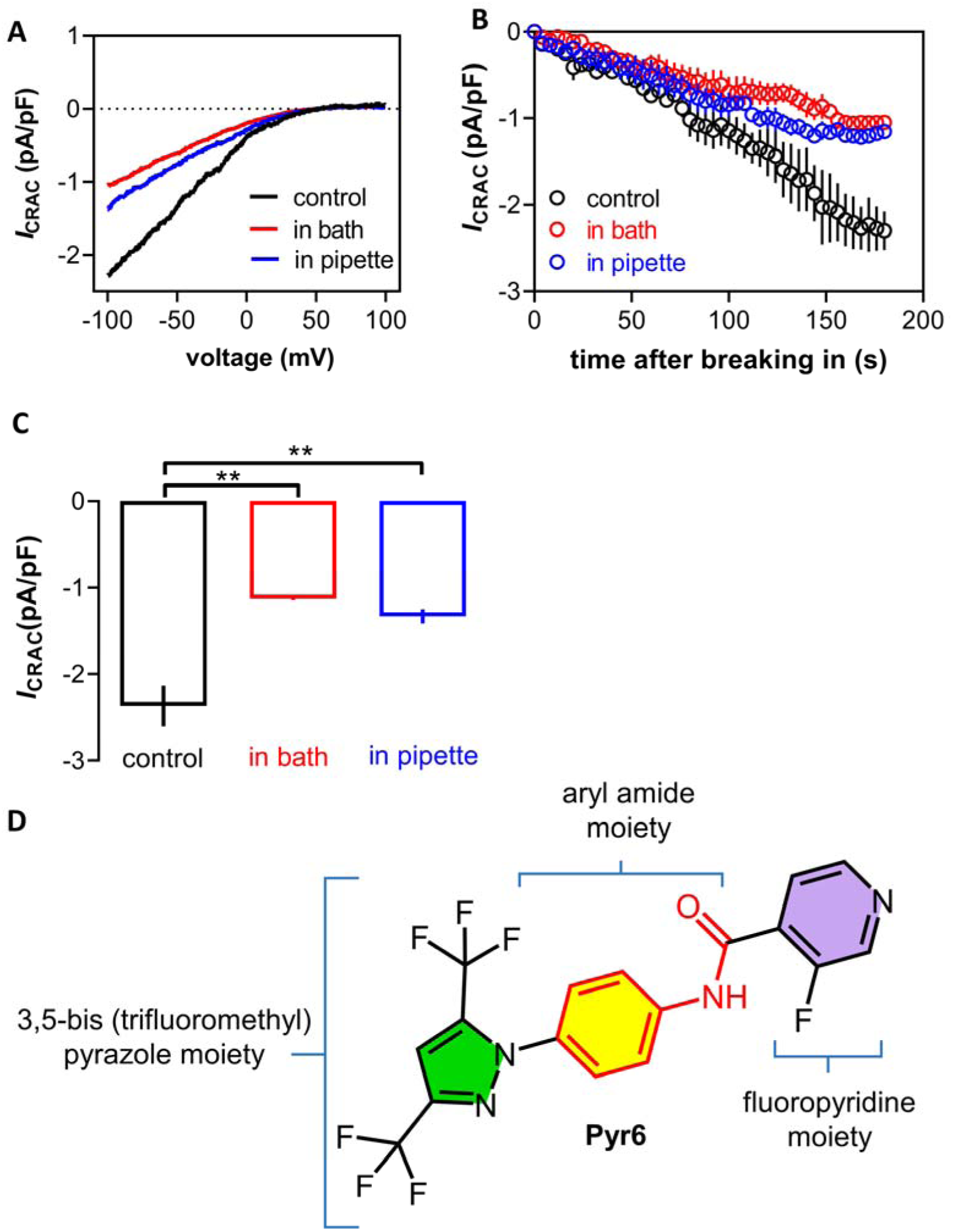
Effects of Pyr6 on CRAC channel function in RBL-1 cells. Prior to recording of the whole-cell current representing CRAC channel function (*I*_CRAC_), RBL-1 cells were pre-treated with Pyr6 (10μM) for ∼20min added to the bath solution or included in the pipette solution whilst control cells were treated with DMSO only. **A**) Typical current-voltage (I-V) relationships of ICRAC (pA) evoked by voltage ramps from −100mv to +100mv, normalized to cell size (pF) under different experimental conditions. **B)** Representative peak *I*_CRAC_ size (as current density, pA/pF) vs time plots (each data point as mean ± SEM) from whole-cell recording of at least four different cells (n =4) in each condition are shown. **C)** Bar graphs (mean ± S.E.M) showing current densities at −80 mV holding potential under each condition. The statistical comparisons with respect to the control were done using Student’s unpaired t-test, **P<0.01. **D**) 2D chemical structure of Pyr6, highlighting different sub-structural features.

However, such subtle change (which is likely to be a case of bioisosteric replacement) seems to have conferred significantly higher selectivity and potency for CRAC channels relative to TRPC3 [22]. In our whole cell recording with RBL-1 cells, Pyr6 at 10μM behaved in stark contrast with that of BTP2 meaning significant suppression of *I*_CRAC_ was observed when it was present either in the extracellular (bath) solution or intracellular (pipette) solution (**Fig 2**).

GSK-7975A is a member of the N-pyrazole carboxamide chemical series developed by GlaxoSmithKline (GSK patent WO/2010/122089) and is also well-recognised as an effective inhibitor of CRAC channels [3]. A previous study confirmed that this compound does not interfere with Stim-Stim or Stim-Orai interaction, instead behaving like an allosteric blocker of the Orai pore with its affinity influenced by the geometry of the selectivity filter [12]. From 2D chemical structural point of view, GSK-7975A appears to be distinct from the BTP2 and Pyr6 (**Fig 3**). However, intriguingly when we tested this molecule against *I*_CRAC_ in whole cell recording of RBL-1 cells, it (10μM) effectively phenocopied BTP2 in significantly suppressing *I*_CRAC_ only when present at the extracellular side (**Fig 3**).

**Fig 3.**
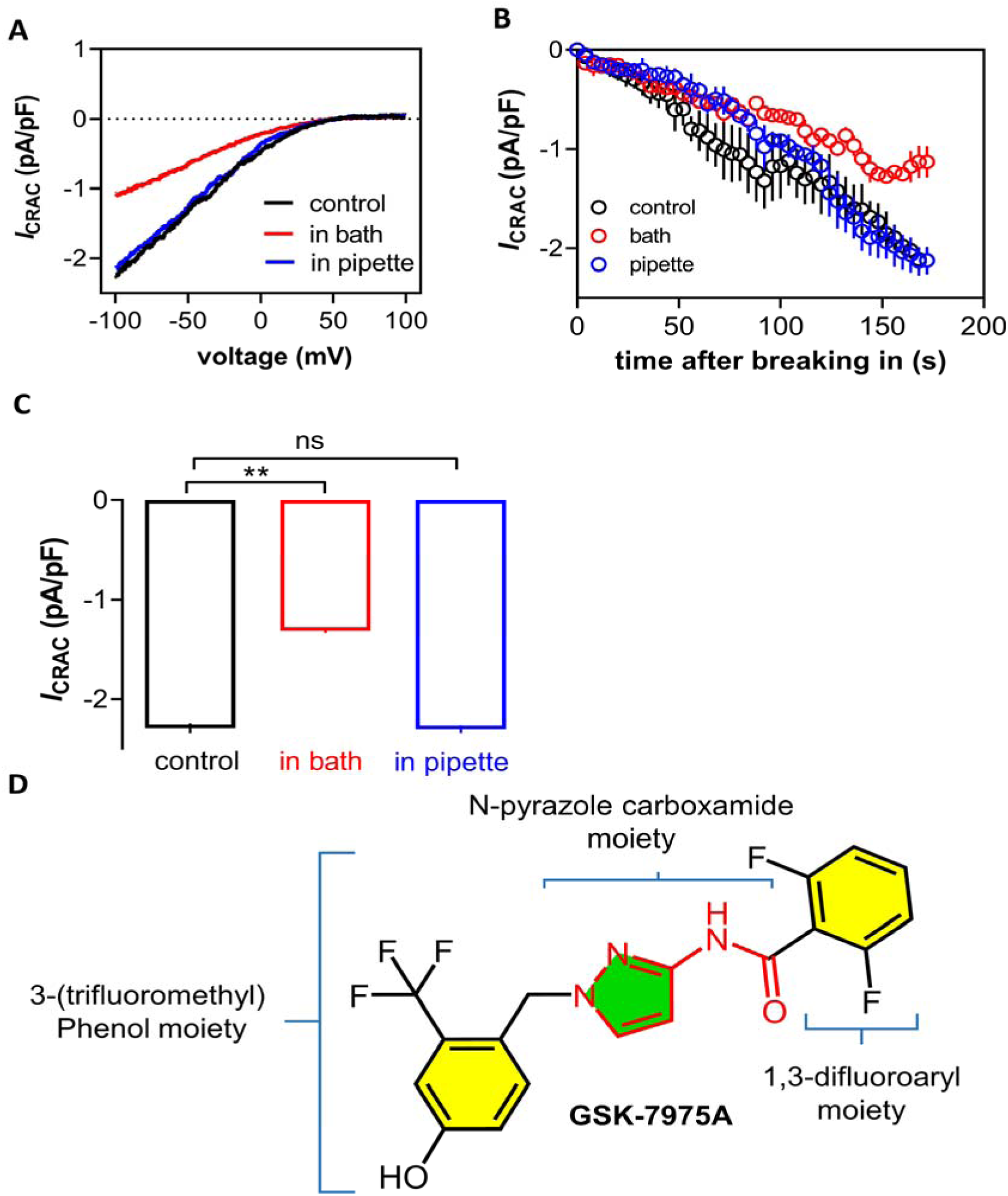
Effects of GSK-7975A on CRAC channel function in from RBL-1 cells. Prior to recording of the whole-cell current representing CRAC channel function (*I*_CRAC_), RBL-1 cells were pre-treated with GSK-7975A (10μM) for ∼20min added to the bath solution or included in the pipette solution whilst control cells were treated with DMSO only. **A**) Typical current-voltage (I-V) relationships of ICRAC (pA) evoked by voltage ramps from −100mv to +100mv, normalized to cell size (pF) under different experimental conditions. **B)** Representative peak *I*_CRAC_ size (as current density) vs time plots (each data point as mean ± SEM) from whole-cell recording of at least four different cells (n=4) in each condition are shown. **C)** Bar graphs (mean ± S.E.M) showing current densities at −80 mV holding potential under each condition. The statistical comparisons with respect to the control were done using Student’s unpaired t-test, **P<0.01. **D**) 2D chemical structure of GSK-7975A, highlighting different sub-structural features.

Synta66 is a selective and efficacious inhibitor of SOCE (and thus CRAC channels) developed by Synta Pharmaceuticals (now merged into Madrigal Pharmaceuticals) [23]. This molecule did not affect clustering of Stim proteins when studied in vascular smooth muscle cells [24] and a recent study [13] has suggested it to be an allosteric pore blocker of Orai1, similar to GSK −7975. From 2D chemical structural point of view, it very closely mimics Pyr6 with the only difference being the 3,5-bistrifluoromethyl pyrazole ring replaced by a 2,5-dimethoxy benzene ring (**Fig 4**).

**Fig 4.**
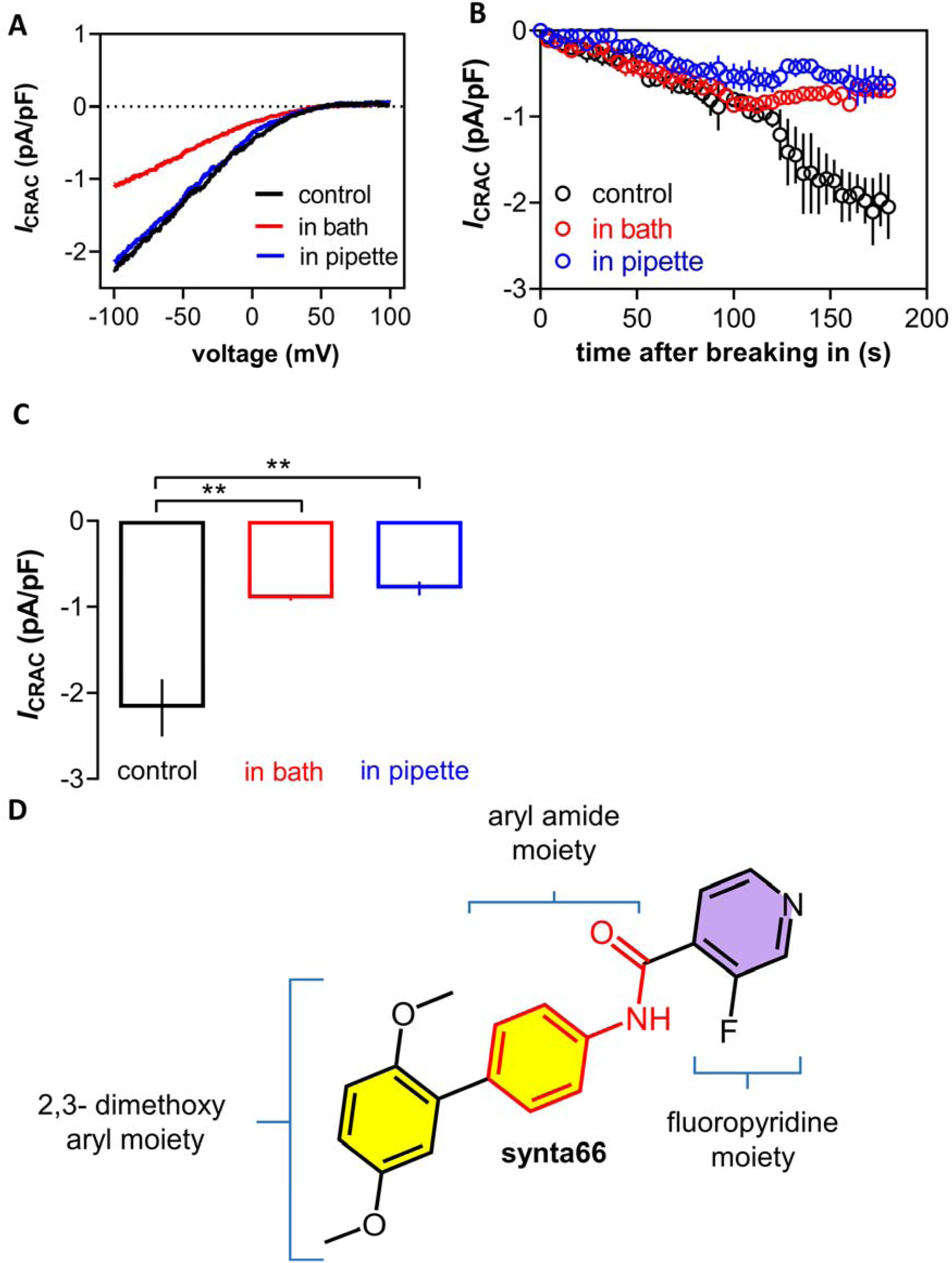
Effects of Synta66 on CRAC channel function in RBL-1 cells. Prior to recording of the whole-cell current representing CRAC channel function (*I*_CRAC_), RBL-1 cells were pre-treated with BTP2 (10μM) for ∼20min added to the bath solution or included in the pipette solution whilst control cells were treated with DMSO only. **A**) Typical current-voltage (I-V) relationships of ICRAC (pA) evoked by voltage ramps from −100mv to +100mv, normalized to cell size (pF) under different experimental conditions. **B)** Representative peak *I*_CRAC_ size (as current density) vs time plots (each data point as mean ± SEM) from whole-cell recording of at least four different cells (n=4) in each condition are shown. **C)** Bar graphs (mean ± S.E.M) showing current densities at −80 mV holding potential under each condition. The statistical comparisons with respect to the control were done using Student’s unpaired t-test, **P<0.01. **D**) 2D chemical structure of Synta66, highlighting different sub-structural features.

Quite intriguingly, when evaluated against *I*_CRAC_ in our whole cell recordings of RBL-1 cells, Synta 66 also functionally mimicked Pyr6 meaning it too was equally able to suppress the peak *I*_CRAC_ size from either side of the plasma membrane at 10μM concentration (**Fig 4**). Thus, in our experimental testing of four popular CRAC channel inhibitors against *I*_CRAC_ representing native CRAC channels in RBL-1 cells, we found BTP2 and GSK-7975A to be acting only extracellularly whilst Pyr6 and Synta66 being able to suppress CRAC channel function when added to either side of the plasma membrane. Given their comparable physicochemical properties (**S1 Table**), we reasoned that the observed differences in the mode of CRAC channel inhibition by these molecules were unlikely due to any possible difference in their cell permeability. Based on these findings, we next sought out to predict plausible site(s) of interaction of these molecules using *in silico* modelling and docking.

### Putative binding mode of selected CRAC inhibitors on rat Orai1 hexamer

For unbiased prediction of the putative binding site(s) and binding mode(s) of the selected CRAC inhibitors evaluated electrophysiologically in our study, we used an *in silico* protocol that we published previously[19, 20]. Since the primary cell line (RBL-1 cells) we used in our electrophysiology is of rat origin, we first made a homology model of rat Orai1 (rOrai1) hexamer based on published crystal structure of Drosophila Orai (PDB id: 4HKR). We considered a hexamer of rOrai1 because of the prevailing broad consensus across the field that the functional CRAC channels in physiological setting comprise of hexameric Orai proteins alongside the STIM proteins[1].

The most frequently-visited site for BTP2 in our *in silico* docking protocol was between the pore-forming M1 helices of two consecutive rat Orai1 subunits (**Fig 5**).

**Fig 5.**
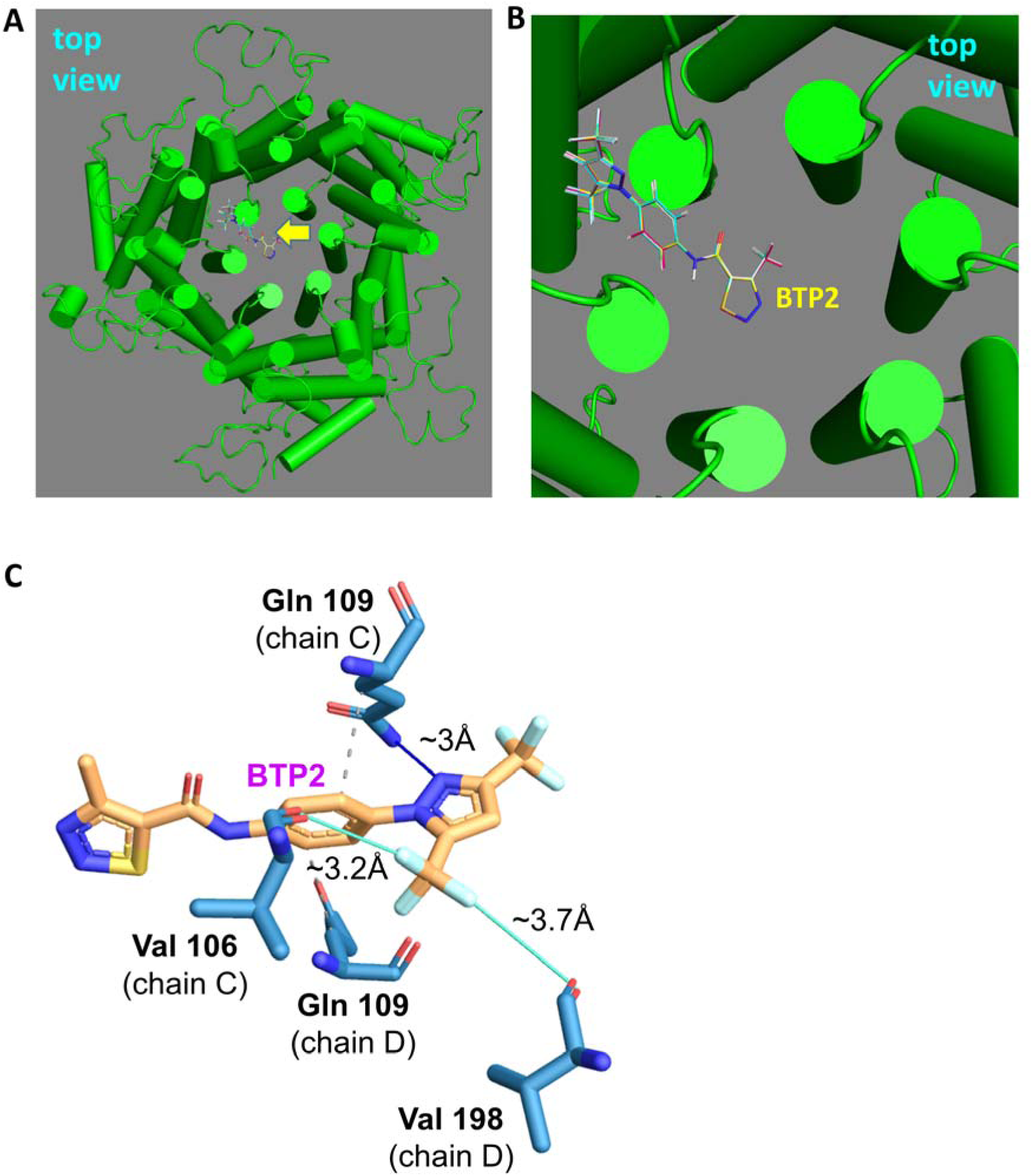
*In silico* prediction of probable binding site and bind mode of BTP2 on modelled rOrai1 hexamer. **A)** Top view from the extracellular side showing the best-ranked and most reproducible binding poses (shown as line representation) of BTP2 from 10 independent docking runs using GOLD suite) on modelled rOrai1 hexamer (coloured green). Position of docked ligand poses are shown with yellow arrow. **B)** close-up view of the image shown in panel A. **C**) 3D-ligand interaction profile of the typical best pose of BTP2 generated through PLIP server. The ligand and the proximal residues are shown as sticks. The figure was obtained from the PLIP server ((https://plip-tool.biotec.tu-dresden.de/plip-web/plip/index).

In this mode, the methyl thiadiazole moiety of BTP2 appears to projects towards the pore lumen whilst the intermediary aryl amide group seems to span the two M1 helices from two Orai1 subunits and the 3,5-bis (trifluoromethyl) pyrazole moiety faces towards the M3 helix. Analysis of the representative docked complex through PLIP revealed a hydrogen bonding interaction between one of the nitrogen atoms of the pyrazole ring and Gln109 (Gln108 for hOrai1). The intermediary aryl group appears to have hydrophobic interaction with Gln109 of both the subunits. One of the trifluoromethyl group engages into halogen bonding with Val106 (Val105 for hOrai1) of one subunit and Val198 (Val 197 for hOrai1) of another subunit.

Chemically speaking, Pyr6 is effectively a hybrid of BTP2 and Synta 66 where the 2,3-dimethoxy aryl moiety of the latter has been replaced by the 3,5-bis (trifluoromethyl) pyrazole moiety of the former (**Fig 2**). Similar to Synta 66 but unlike BTP2, this molecule could significantly suppress *I*_CRAC_ from when present in either the extracellular or the intracellular solution. Therefore, we considered predicting both extracellular and intracellular sites for its interaction with the rOrai1 hexamer in our *in silico* docking approach.

The most frequently-obtained best ranked docked pose of Pyr6 at the extracellular of rOrai1 hexamer was similar to that of BTP2 where the molecule mainly traverses the pore-forming M1 helices of two adjacent Orai1 monomers (**Figs 6A-6C**).

**Fig 6.**
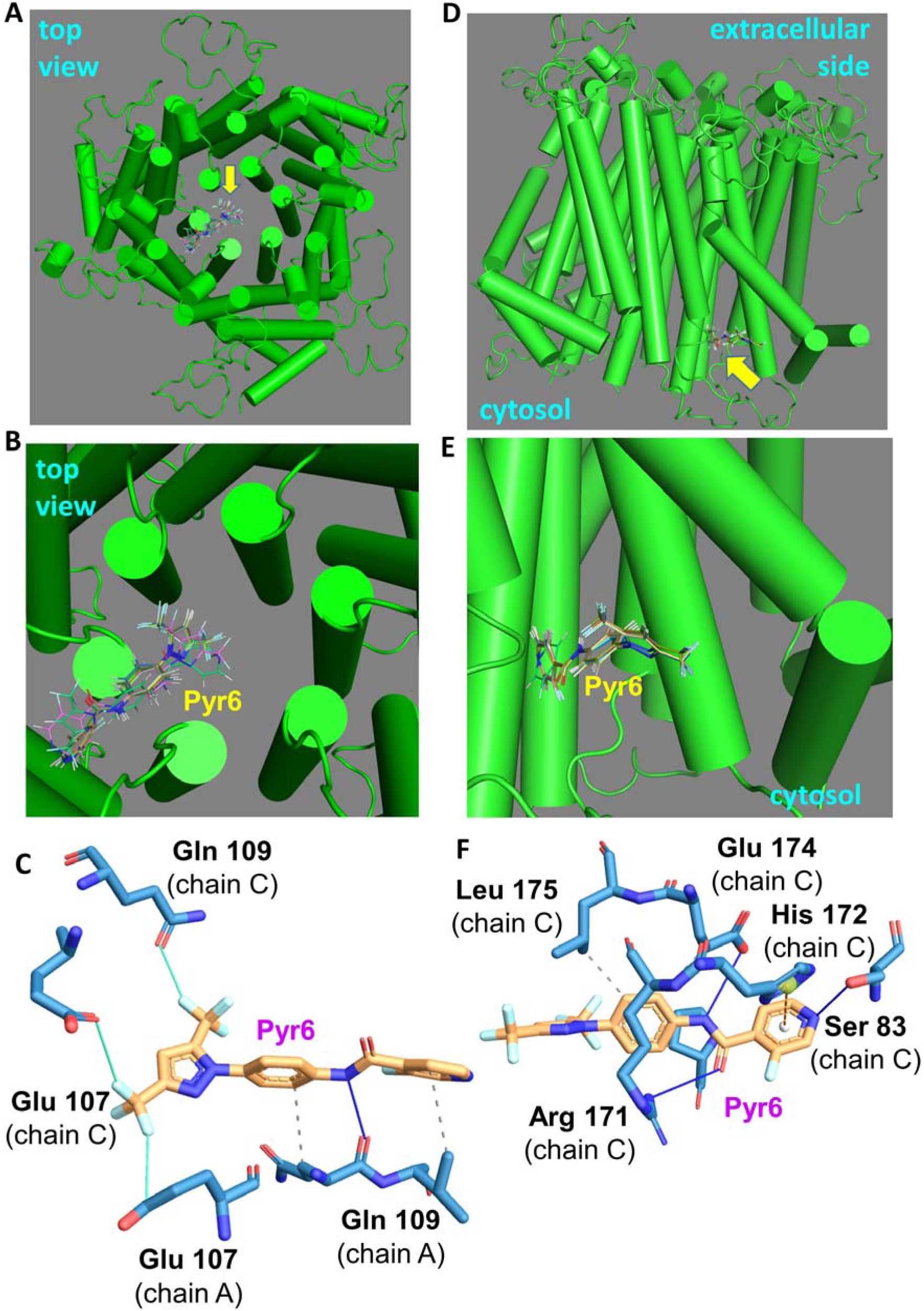
*In silico* prediction of probable binding site and bind mode of Pyr6 on modelled rOrai1 hexamer. **A)** Top view from the extracellular side showing the best-ranked and most reproducible binding poses (shown as line representation) of Pyr6 from 10 independent docking runs using GOLD suite) on modelled rOrai1 hexamer (coloured green). Position of docked ligand poses are shown with yellow arrow. **B)** close-up view of the image shown in panel A. **C**) 3D-ligand interaction profile of the typical best pose of Pyr6 generated through PLIP server. The ligand and the proximal residues are shown as sticks. The figure was obtained from the PLIP server. **D**) Tilted-side view or rOrai1 hexamer model, showing the best-ranked and most reproducible binding poses of Pyr6 (shown as line representation) of Pyr6 from 10 independent docking runs using GOLD suite) facing towards the cytosol. Position of docked ligand poses are shown with yellow arrow. **E)** close-up view of the image shown in panel D. **F**) 3D-ligand interaction profile of the typical best pose of Pyr6 towards the cytosol. The ligand and the proximal residues are shown as sticks. The figure was obtained from the PLIP server.

In this mode, the trifluoromethyl groups appear to form halogen bonds [25] with Glu 107 (Glu 106 for hOrai1) of one Orai1 subunit and with Glu 106 (Glu106 for hOrai1) and Gln 109 (Gln 108 for hOrai1) from the other subunit. There is a hydrogen bonding interaction between the backbone carbonyl of Gln 109 (Gln108 for hOrai1) of one subunit and the nitrogen atom of the intermediary aryl amide moiety of Pyr6. Gln 109(Gln108 for hOrai1) of one Orai1 subunit appears to form hydrophobic interactions with the intermediary aryl moiety and the fluoropyridine moiety of Pyr6 (**Figs 6A-6C**). The most frequently-visited intracellular site for Pyr6 in our *in silico* docking protocol was a pocket within the vicinity of the lower ends of the pore-forming M1 helix and M3 helix from the same rOrai1 subunit (**Figs 6D-6F**). This site, though neighbouring, is distinct from that the predicted intracellular binding site for Synta 66 (see later). PLIP-based analysis of the relevant docked Pyr6-rOrai1 hexamer complex suggested that there are π-π stacking occurring between the central aryl amide moiety and Arg 171 (Arg 170 for hOrai1) and Glu 174 (Glu 173 for hOrai1) as well as between the fluoropyridine moiety and Ser 83 (Ser 82 for hOrai1). Additionally, there appears to be a cation-π interaction between His 172 (His 171 for hOrai1) and the fluoropyridine moiety whilst Leu 175(Leu174 for hOrai1) engages into a hydrophobic interaction with the central aryl region of Pyr6 (**Figs 6D-6F**).

GSK7975A with an intermediary pyrazole amide moiety is chemically somewhat distinct from BTP2, Pyr6 and Synta66 that possess an aryl amide moiety flanking the two ends of their structures. Interestingly, in our *in silico* docking, GSK-7975A poses were clustered to two adjacent but distinct binding sites. The predicted site 1 (shown in yellow arrow, **Fig 7**) and the binding mode for GSK-7975A appear to be similar to the three other inhibitors where the trifluoromethyl phenol and the intermediary pyrazole moieties stay in close proximity to the pore-forming M1 helices of two adjacent rat Orai1 subunits.

**Fig 7.**
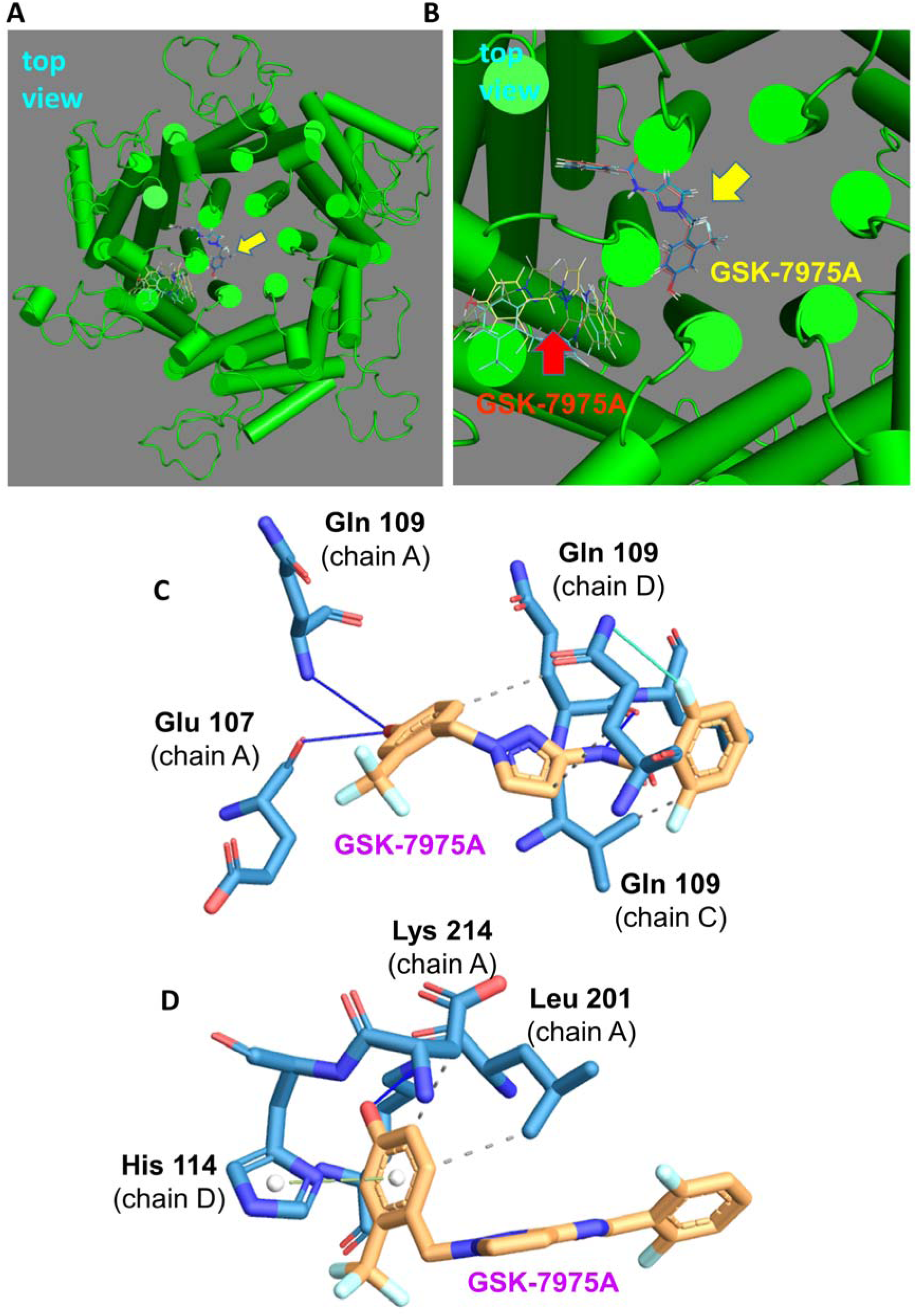
*In silico* prediction of probable binding site and bind mode of GSK-7975A on modelled rOrai1 hexamer. **A)** Top view from the extracellular side showing the best-ranked and most reproducible binding poses (as line representation) of GSK-7975A from 10 independent docking runs on modelled rOrai1 hexamer (coloured green). Position of docked ligand poses are shown with yellow arrow. **B)** closed-up view of the image shown in panel A. The two distinct sites are shown with red and yellow arrow signs. Position of docked ligand poses are shown with yellow arrow. **C**) 3D-ligand interaction profile of the typical best pose of GSK-7975A at site 1 (shown with yellow arrow sign) generated through PLIP server. The ligand and the proximal residues are shown as sticks. The figure was obtained from the PLIP server. **D**) 3D-ligand interaction profile of the typical best pose of GSK-7975A at site 2 (shown with red arrow sign) generated through PLIP server. The ligand and the proximal residues are shown as sticks. The figure was obtained from the PLIP server.

The difluoroaryl moiety at the other end of the molecule juxtaposes the M3 helix of one Orai1 subunit. PLIP-based analysis reveals that the phenolic group of the trifluoromethyl phenol moiety engages into two hydrogen bonding with Gln 109 (Gln 108 for hOrai1) and Glu 107 (Glu 106 for hOrai1) of one Orai1 subunit. Gln 109 (Gln 108 for hOrai1) of a different Orai1 subunit seems to form two hydrogen bonding with one of the pyrazole nitrogen atoms and the adjacent amide nitrogen atom. The difluoroaryl moiety mainly forms hydrophobic interactions with Val 108 (Val 107 for hOrai1), Gln 109 (Gln 108 for hOrai1) and Leu 110 (Leu 109 for hOrai1) of one Orai1 subunit and with Gln 109 (Gln 109 for hOrai1) of another but adjacent Orai1 subunit. Finally, there is a halogen bond formed between the amide nitrogen atom of Gln 109 (Gln 108 for hOrai1) of one Orai1 subunit and one of the fluorine atom of the difluoroaryl moiety of the molecule (**Fig 7**). Unlike three other CRAC inhibitors used in this study, docked poses of GSK-7975A also clustered (with almost similar frequency to site 1) at another site (site 2, shown with red arrow, **Fig 7**) where the entire molecule appears to span the space between the pore-forming M1 helix and the M3 helix of rat Orai1. PLIP-based analysis reveals that the larger part of the molecule in this mode remains solvent exposed (not in explicit interaction with any Orai1 residues but the trifluoromethyl phenyl moiety manifests several interactions: hydrogen bonding of the phenolic group with Lys 214 (Lys 215 and hydrophobic interaction with Leu 201 (Leu 200 for hOrai1) of one Orai1 subunit; π-π stacking between the aryl ring and the imidazole ring of His 114 (His 113 for hOrai1) as well as hydrophobic interaction with Asp 113 (Asp 112 for hOrai1) of a different but neighboring Orai1 subunit (**Fig 7**). It is to be mentioned here that compared to the site 1, poses of GSK-7975A showed more variability in their absolute orientation with respect to the rOrai1 model. Like BTP2, synta66 retains the core aryl amide moiety but this is flanked by a fluoropyridine moiety and a 2,3-dimethoxyaryl moiety (**Fig 4**). Unlike BTP2 but similar to Pyr6, Synta 66 was equally effective in suppressing *I*_CRAC_ when introduced to bath (extracellular) or the pipette (intracellular) solution. We therefore considered both extracellular and intracellular site of interaction of this molecule on the rat Orai 1 hexamer model. The putative extracellular binding site of Synta 66 appears to be analogous to what was observed for BTP2 and Pyr6 (discussed below) meaning it is between the pore-forming M1 helices of two consecutive Orai1 subunits (**Figs 8A-8C**).

**Fig 8.**
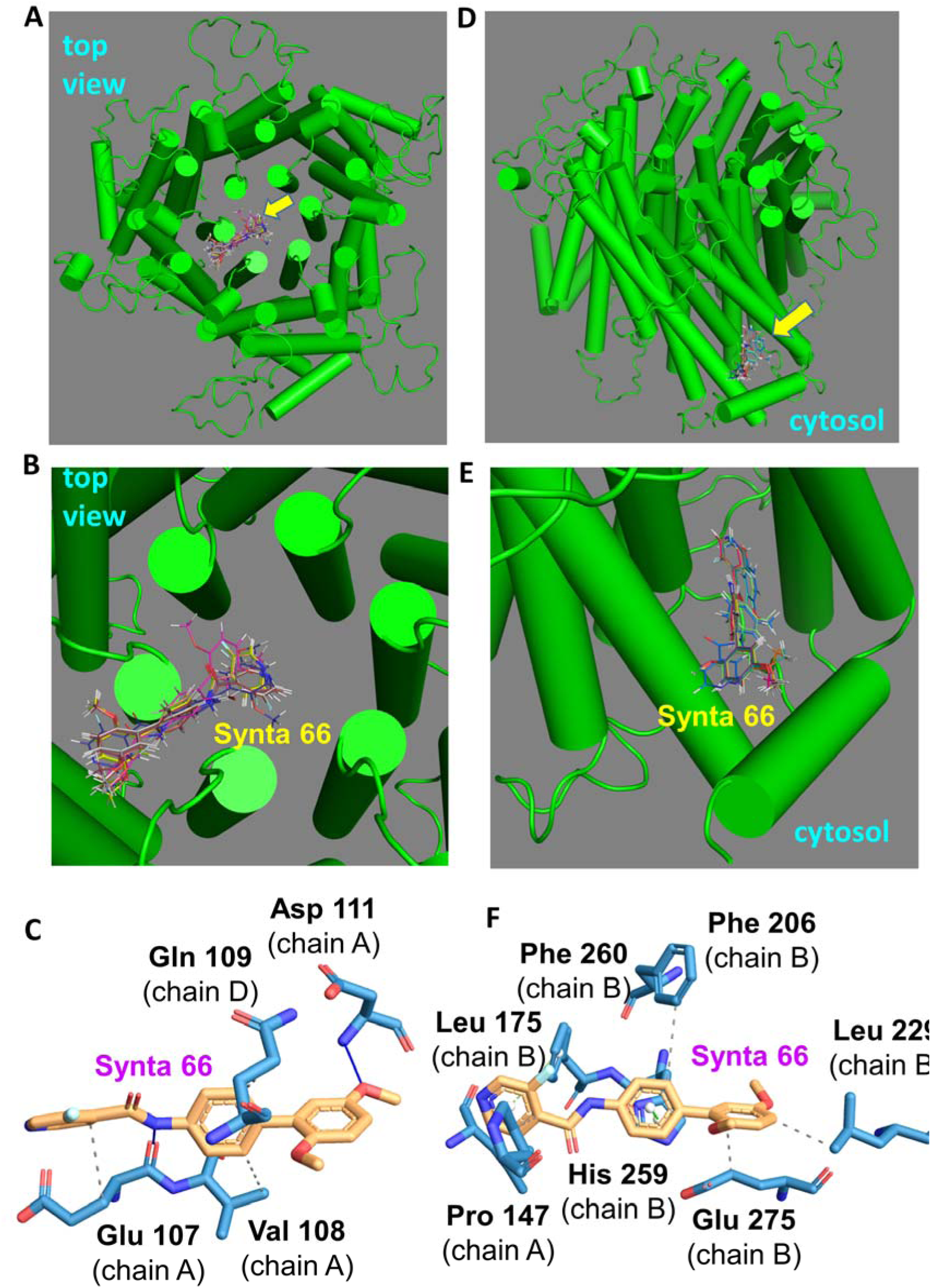
*In silico* prediction of probable binding site and bind mode of Synta66 on modelled rOrai1 hexamer. **A)** Top view from the extracellular side showing the best-ranked and most reproducible binding poses (shown as line representation) of Synta66 from 10 independent docking runs using GOLD suite) on modelled rOrai1 hexamer (coloured green). Position of docked ligand poses are shown with yellow arrow. **B)** close-up view of the image shown in panel A. **C**) 3D-ligand interaction profile of the typical best pose of Synta66 generated through PLIP server. The ligand and the proximal residues are shown as sticks. The figure was obtained from the PLIP server. **D**) Tilted-top view or rOrai1 hexamer model, showing the best-ranked and most reproducible binding poses (shown as line representation) of Synta66 from 10 independent docking runs using GOLD suite) facing towards the cytosol. Position of docked ligand poses are shown with yellow arrow. **E)** close-up view of the image shown in panel D. **F**) 3D-ligand interaction profile of the typical best pose of Synta66 towards the cytosol. The ligand and the proximal residues are shown as sticks. The figure was obtained from the PLIP server.

Analysis of the representative docked complex through PLIP revealed a hydrogen bonding interaction between one of the methoxy groups on the terminal aryl moiety and Asp 111 (Asp 110 for hOrai1). There appears to be another hydrogen bonding interaction between the amide group of the molecule and Glu 107 (Glu 106 for hOrai1). The intermediary aryl group appears to have hydrophobic interaction with Val 108 (Val 107 for hOrai1) whilst similar hydrophobic interaction seems to occur between the terminal fluoropyridine group and Glu 107 (Glu 106 for hOrai1).

The most frequently-visited intracellular site for Synta66 in our *in silico* docking protocol was a pocket within the vicinity of the lower ends of M3 and M4 helices from consecutive Orai1 subunits (**Figs 8D-8F**). PLIP-based analysis suggested that there are π-π stacking occurring between the central aryl moiety and His 259 (His 256 for hOrai1) and between the fluoropyridine moiety and Phe 260 (Phe 267 for hOrai1). Other than these, the recognition of Synta 66 at this cavity is largely governed by hydrophobic interactions involving Pro 147 (Pro 146 for hOrai1), Leu 175 (Leu 174 for hOrai1), Phe 256 (Phe 253 for hOrai1), Glu 275 (Glue 272 for hOrai1) and Leu 279 (Leu 276 for hOrai1).

Of all the residues that seem to be involved in the recognition of these, there are residues that are directly part of the pore-forming M1 helix of Orai1 protein and these include Val 107 (Val 106 for hOrai1), Gln 109 (Gln 108 for hOrai1) and Glu 107 (Glu 106 for hOrai1). Of these, the highly conserved Glu 106 is well known to be critical for governing Ca^2+^ selectivity, pore diameter, pH regulation and Ca^2+^-dependent inhibition [1, 2]. Intriguing, a previous study by Romanin and colleagues showed that mutation of this Glu to Asp (Glu 106 to Asp 106 for hOrai1), besides reducing Ca^2+^ selectivity, also abolished the inhibition by GSK-7975A [12]. In this context, our finding that GSK-7975A only inhibits CRAC current from the extracellular end is entirely plausible and our proposed ‘site I’ binding mode for GSK-7975A strongly complies with their finding but this also suggests that our predicted ‘site II’ binding mode is likely to be unrealistic for this molecule.

Like GSK-7975A, BTP2 in our hand also was only effective extracellularly in inhibiting CRAC channel. Interestingly this agrees in an early finding in Jurkat T cells where the authors suggested extracellular mode of action of BTP2 [14]. In our in silico blind docking, BTP2 does not seem to interact with the conserved Glu 107 (Glu 106 for hOrai1) directly but it does with the adjacent Val 106 (Val 105 for hOrai1) whilst among few other interacting residues, Gln 109 (Gln 108 for hOrai1) is also in close proximity to Glu 107 – in the loop just above the pore-forming M1 helix whilst Val 198 (Val 197 for hOrai1) is located in M3 helix which is just behind the pore-forming M1 helix. Such binding site of BTP2 will be compatible with its extracellular mode of action as we and others have found [14].

Despite structural similarity with BTP2 and GSK-7975A, the other two CRAC inhibitors namely Pyr6 and Synta 66 were found to be effective given either extracellularly or intracellularly. Their extracellular binding site involves the conserved Glu 107 (Glu 106 for hOrai1) and few in its neighbourhood including Gln 109 (Gln 108 for hOrai1) as observed for BTP2 and GSK-7975A compounds. Besides this site, we also suggest that Pyr6 and Synta 66 may also act via intracellular sites which may also be plausible as suggested previously [4]. Another aspect that deserves our attention is the fact that for each CRAC inhibitor, we only considered 1:1 stoichiometry during our in silico blind docking in agreement with previous studies [12] involving some of the CRAC inhibitors (GSK-7975A and Synta66) that showed a Hill coefficient of ∼1. We nevertheless cannot rule out multiple occupancies of these inhibitors for which solving inhibitor-bound Orai structure will be essential.

## Conclusion

The overall aim of this study was to determine, primarily using whole-cell patch clamp recording of *I*_CRAC_ from RBL-1 cells, the mode of action of four widely-used CRAC channel inhibitors being extra- or intracellular or both. Despite having comparable physicochemical properties and some chemical similarities, we observed that these molecules differ in their mode of action – two (BTP2 and GSK-7975A) being effective only extracellularly whilst the others can inhibit *I*_CRAC_ when added to the either side of the membrane. Our *in silico* binding site prediction for these molecules against modelled rOrai1 hexamer suggest sites that either directly involve or proximal to the pore-forming M1 helix from extracellular or intracellular end relevant for specific blocker. If these sites are valid, this would then imply that these CRAC channel inhibitors cause some allosteric distortion of the Orai1 pore geometry and/or affect the rotation of M1 helix and subsequent pore dilatation. The resultant inappropriate pore geometry may not only directly impede Ca^2+^ permeation but also may affect the channel gating[26]. Our findings and hypotheses agree well with suggestions from few studies[12–14]. We also provide here with some amino acids of Orai1 proteins that are likely to be involved in recognition of these popular CRAC inhibitors. Also, for inhibitors that seem directly engage the conserved Glu 107 (Glu 106 for hOrai1), we anticipate that these ligands may also interact with Ca^2+^ which requires further investigation involving either solving the inhibitor-bound Orai1 structure and/or with molecular dynamics simulation. Finally for two CRAC inhibitors namely Synta66 and Pyr6, we suggest potential two sites of action.

In this work, we only relied on endogenous CRAC channels in a popular mast cell line namely RBL-1 cells which is likely to contain all three Orai isoforms. Future studies may consider site directed mutagenesis of specific Orai1 and functional re-evaluation of these inhibitors and also evaluating the same for Orai2 and Orai3 isoforms expressed in a Orai-null background as used before [27]. This will be important, given some difference observed in pharmacology across Orai isoforms [28]. Future work should also involve cells with overexpressed CRAC machineries such as Stim and/or Orai proteins to check whether or not the observed behaviour of these CRAC inhibitors against endogenous CRAC channels also hold for overexpressed system.

## Data availability

The data presented in this study can be obtained from the corresponding author upon reasonable request.

## Acknowledgements

This study was supported by a grant from The Scientific and Technological Research Council of Turkey (TUBITAK-1059B141800131). Sinayat Mahzabeen was funded by a PhD studentship from the Cambridge Trust. The funders had no role in study design, data collection and analysis, decision to publish, or preparation of the manuscript

## Authors’ contributions

TR conceived the project. YG performed all the electrophysiological experiments; SM and JG carried out all the *in silico* modelling. YG and TR wrote the manuscript with contributions from SM, JG, MH and NY. TR overall supervised the project.

## Supporting information

**S1 Fig. Cartoon representation of the rat Orai 1 single subunit model from AlphaFold. The model was retrieved from AlphaFold** (https://alphafold.ebi.ac.uk/entry/Q5M848) and the disordered region is highlighted in red colour. The figure was generated in PyMOL Molecular Graphics System (version 1.8.2.0 Open-Source, Schrödinger, LLC).

**S1 Table. Predicted physicochemical properties of the CRAC channel inhibitors used in the study**

*MW, Molecular Weight; XlogP3, predicted octanol-water partition coefficient; TPSA, total polar surface area. Values are predicted and taken from PubChem) https://pubchem.ncbi.nlm.nih.gov/).

