## Supplementary figures and images for "Re-evaluation of some popular CRAC channel inhibitors – structurally similar but mechanistically different?"

### Supplemental Figure 1

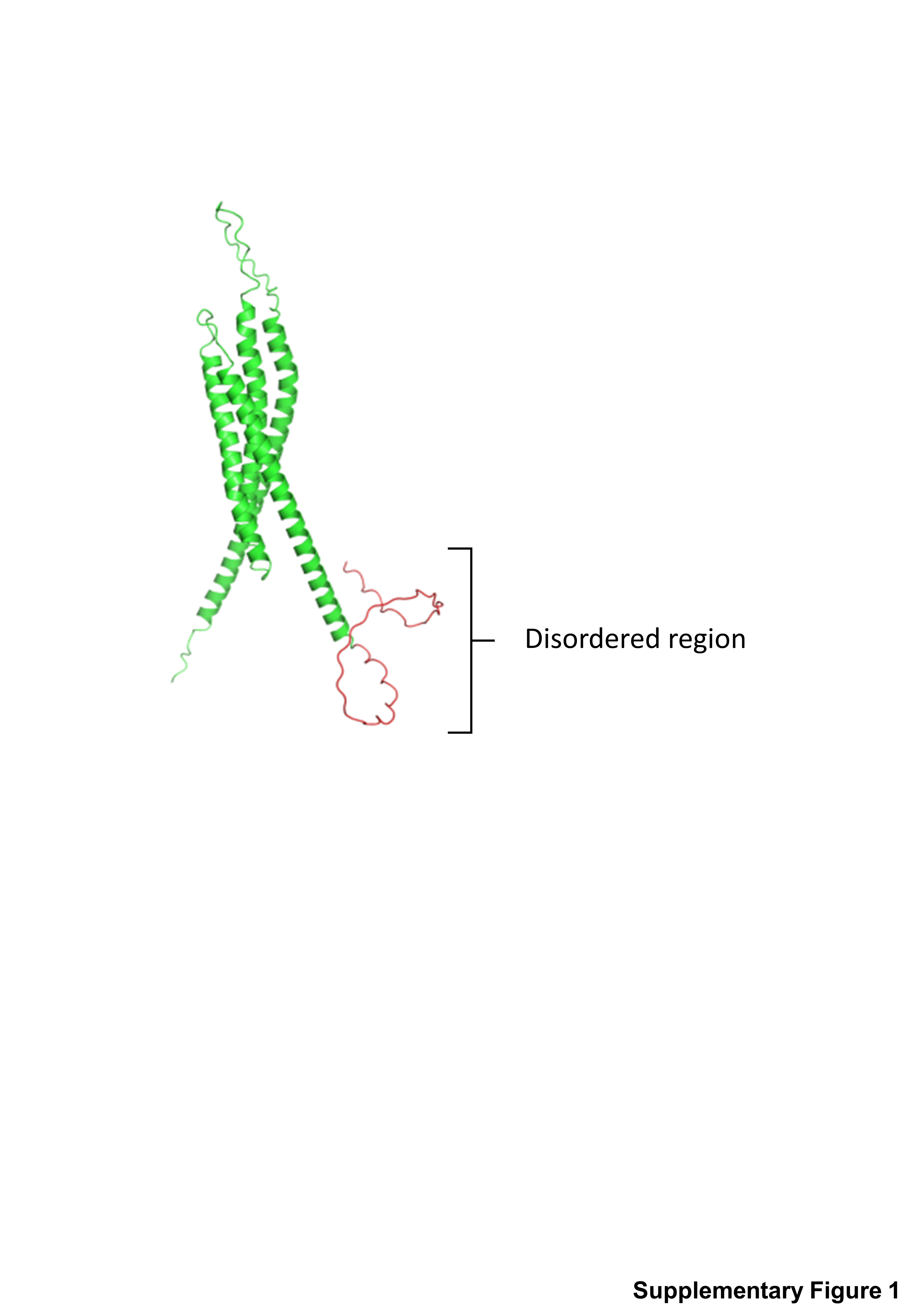
